# gp120 envelope glycoproteins of HIV-1 Group M Subtype A and Subtype B differentially affect gene expression in human vascular endothelial cells

**DOI:** 10.1101/2023.01.03.522636

**Authors:** Andrew J. Suh, Dante I. Suzuki, Sergiy G. Gychka, Tinatin I. Brelidze, Yuichiro J. Suzuki

## Abstract

Cardiovascular complications are seen among human immunodeficiency virus (HIV)-positive individuals who can now survive longer due to successful antiretroviral therapies. Among them, pulmonary arterial hypertension (PAH) is a fatal disease characterized by increased blood pressure in the lung circulation due to vasoconstriction and vascular wall remodeling, resulting in the overworking of the heart. The prevalence of PAH in the HIV-positive population is dramatically higher than that in the general population. While HIV-1 Group M Subtype B is the most prevalent subtype in western countries, the majority of HIV-1 infections in eastern Africa and former Soviet Union countries are caused by Subtype A. Research on the mechanism of vascular complications in the HIV-positive population, especially in the context of subtype differences, however, has not been rigorous. Much of the research on HIV has focused on Subtype B and information on the molecular mechanisms of Subtype A is non-existent. The lack of such knowledge results in health disparities in the development of therapeutic strategies to prevent/treat HIV complications. The present study examined the effects of HIV-1 viral fusion protein gp120 of Subtypes A and B on cultured human pulmonary artery endothelial cells by performing protein arrays. We found that the gene expression changes caused by the gp120s of Subtypes A and B are different. Specifically, Subtype A is a more potent downregulator of perostasin, matrix metalloproteinase-2 (MMP-2), and ErbB/Her3 than Subtype B, while Subtype B is more effective in downregulating monocyte chemotactic protein-2 (MCP-2/CCL8), MCP-3 (CCL7), and thymus- and activation-regulated chemokine (TARC/CCL17) proteins. This is the first report of gp120 proteins affecting host cells in an HIV subtype-specific manner, opening up the possibility that vascular complications may occur differently in HIV patients throughout the world.

## Introduction

The increased incidence of cardiovascular complications is seen among human immunodeficiency virus (HIV)-positive individuals who can now survive longer because of successful antiretroviral therapies (ART). Among them, pulmonary arterial hypertension (PAH) is a serious and fatal disease characterized by increased blood pressure in lung circulation due to vasoconstriction and vascular wall remodeling, resulting in the overworking of the heart. HIV-associated PAH occurs in approximately one out of 200 HIV-infected patients (Mehta, 2000; Sitbon, 2008; Isasti, 2013; Pellicelli, 2001), which is dramatically higher than the prevalence of PAH in individuals without HIV (Bigna, 2015). A large prospective study of 7,648 patients with HIV after the availability of potent ART showed a prevalence of right-heart catheterization-confirmed HIV-associated PAH of 0.5% (100 – 1,000 times greater than the prevalence in individuals without HIV infection). These findings are similar to those of studies performed before the development of effective ART (Sitbon et al., 2008). Thus, ART does not seem to have altered the prevalence of HIV-PAH, suggesting the direct role of HIV components in promoting PAH.

Group M Subtype B is the most prevalent subtype of HIV-1 in western countries, where most of the research data on HIV have been generated. By contrast, the majority of HIV-1 infections in eastern African and former Soviet Union countries are caused by Group M Subtype A (Nikolopoulos, 2016; Saad, 2006). However, research on the HIV-1 subtype specificity has not been rigorous, and the information available in the scientific literature may not necessarily apply to the HIV infection in many countries, possibly generating disparity between western countries and low-income countries in developing appropriate therapeutic strategies.

The viral fusion protein of HIV-1, the envelope glycoprotein gp160, consists of a complex of gp120 and gp41 (Checkley et al., 2011). gp120 contains the binding site to the target host cell receptor and is cleaved off from gp160 by cellular proteases including furin (Hallenberger et al., 1992). In the absence of the rest of the viral components, gp120 has also been shown to activate cell signaling pathways, including chemokine receptors, protein kinase C, mitogen-activated protein kinases, and reactive oxygen species in human vascular smooth muscle cells (Schecter et al., 2001). In human pulmonary artery smooth muscle cells, gp120 was found to increase intracellular calcium and induce cell growth (Amsellem et al., 2014). These effects of gp120 were inhibited by an inhibitor of CCR5, a coreceptor for cellular HIV entry. In human lung microvascular endothelial cells, gp120 promoted apoptosis and the secretion of endothelin-1, which mediates the development of PAH (Kanmogne et al., 2005).

These previous studies of the effects of vascular cells used gp120 of HIV-1 Group M Subtype B only. Despite many individuals in the world being infected with HIV-1 Group M Subtype A, information about the cellular effects of gp120 of Subtype A is completely absent. Thus, in the present study, we asked whether gp120 proteins of HIV-1 Group M Subtype A and Subtype B may exhibit different effects on cultured human pulmonary artery endothelial cells.

## Materials and Methods

### Protein sequence analysis

Protein amino acid sequences for gp120 of Subtype A (accession number AAT67478.1) and gp120 of Subtype B (accession number AAA44191.1) were aligned using Clustal Omega multiple sequence alignment tool provided by the European Bioinformatics Institute (Madeira et al., 2022). The percent of identical amino acids was calculated using the equation, (number of identical amino acids) / [(number of identical amino acids) + (number of nonidentical amino acids)].

### Cell culture

Human pulmonary artery endothelial cells (Catalog number C-12241) were purchased from PromoCell GmbH (Heidelberg, Germany). Cells were cultured in Endothelial Cell Growth Medium (PromoCell, Catalog number C-22010) in accordance with the manufacturer’s instructions in 5% CO_2_ at 37°C. Cells at passages 3-6 were placed in low fetal bovine serum (0.4%)-containing medium before the treatment, as routinely performed in experiments on cell signaling [Shults et al., 2018].

Cells were treated with either the recombinant gp120 of HIV-1 Group M Subunit A (Catalog number 40403-V08H, Sino Biological, Inc., Beijing, China), gp120 of HIV-2 Group M Subunit B (Catalog number 40404-V08H, Sino Biological), S1 of SARS CoV-2 spike protein (Catalog number 40591-V08H, Sino Biological), or S1 of the omicron variant SARS CoV-2 spike protein (Catalog number 40591-V08H41, Sino Biological). Characteristics of these recombinant proteins are summarized in Table 1.

**Table 1:**
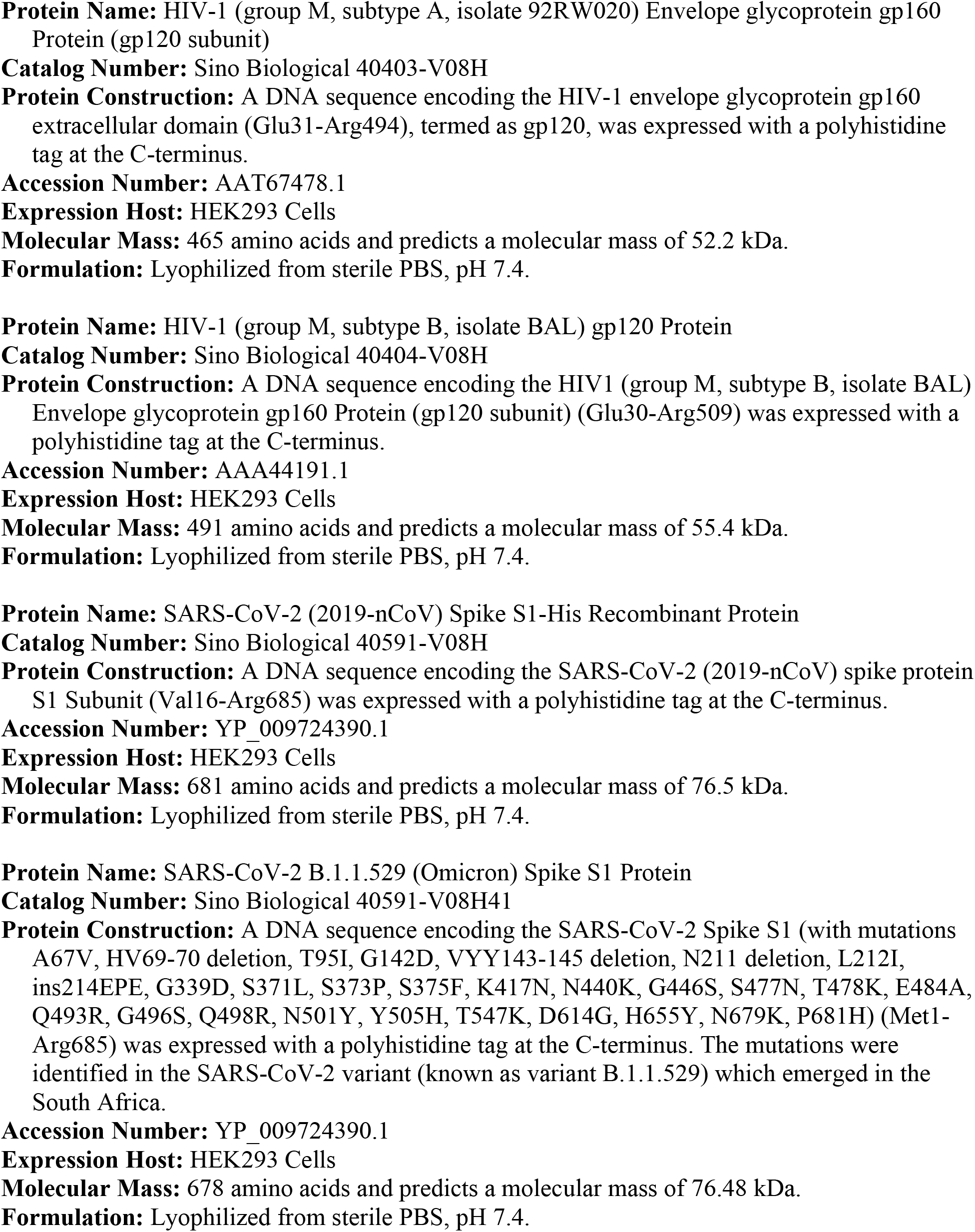
Characteristics of recombinant proteins used in this study

### Protein Array

In this study, Proteome Profiler Array Human XL Oncology Array (Catalog Number ARY026) and Human XL Cytokine Array (Catalog Number ARY022B) from R&D Systems, Inc., Minneapolis, MN, USA) were used. Cell lysates were applied to the protein arrays to detect interactions. To prepare cell lysates, cells were washed in phosphate buffered saline and solubilized with Lysis Buffer 17 (Catalog Number 895943) supplemented with leupeptin (10 μg/ml) and aprotinin (10 μg/ml) by vortexing and gentle rocking at 4°C for 30 min. Samples were then centrifuged at 14,000*g* for 10 min at 4°C, supernatants were collected, and protein concentrations were determined using Bio-Rad Protein Assay (Bio-Rad, Hercules, CA, USA).

Arrays were performed in accordance with the manufacturer’s instructions. Briefly, each membrane was incubated with Array Buffer 6 for 1 hour at room temperature on a rocking platform shaker. Solutions were then replaced with appropriate Array Buffer solutions containing cell lysates (100 μg protein) and incubated overnight at 4°C on a rocking platform shaker. Membranes were washed 3 times with Wash Buffer and incubated in the Detection Antibody solution for 1 hour at room temperature on a rocking platform shaker. Membranes were again washed 3 times with Wash Buffer and incubated with IRDye 800CW Streptavidin (1:2000; LI-COR, Lincoln, NE, USA) for 30 min at room temperature. Lastly, membranes were washed 3 times with Wash Buffer and signals were obtained by using the Odyssey Infrared Imaging System (LI-COR).

### Statistical Analysis

Means and standard errors of mean (SEM) were computed. Two groups were compared by a two-tailed Student’s *t* test, and differences between more than two groups were determined by the analysis of variance (ANOVA). *P* < 0.05 was defined to be statistically significant.

## Results

We first analyzed the sequences of gp160’s from HIV-1 Group M Subtypes A and B. Amino acid sequences of gp160 proteins of HIV-1 Subtype A (Accession # AAT67478.1) and Subtype B (Accession # AAA44191.1) are shown in Fig. 1. The furin cleavage sequence Arg-Glu-Lys-Arg (Pasquato et al., 2007) where gp120 and gp41 become separated are shown in bold. The amino acids that are identical between Subtype A and Subtype B are shown in yellow, and strongly and weakly similar amino acids between the two proteins are shown in green and purple, respectively. We calculated that the amino acid sequences of these two proteins share 75.8% identity. The first amino acids of the recombinant gp120 proteins used in this study are shown in bold red letters and the last amino acids in bold blue. These gp120 proteins share only 70.6% identical amino acids. Thus, we hypothesized that the two proteins may affect cells differently.

**Fig. 1:**
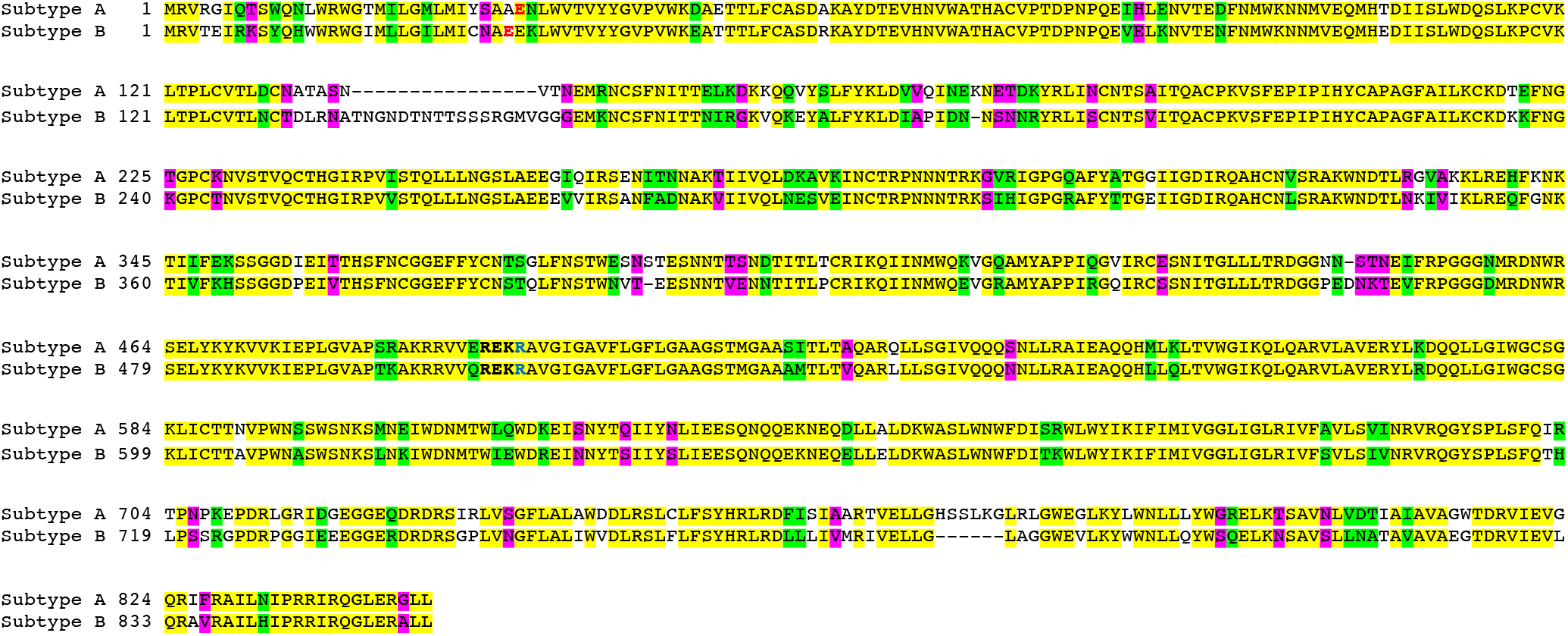
Amino acid sequence alignment of gp160 Subtype A and Subtype B. Amino acid sequence alignment of gp160 proteins of HIV-1 Group M Subtype A (Accession# AAT67478.1) in black and Subtype B (AAA44191.1). The furin cleavage sequence is shown in bold (Pasquato et al., 2007). Amino acids that are identical between Subtype A and Subtype B are highlighted in yellow, and strongly and weakly similar amino acids are highlighted in green and purple, respectively, in accordance with the definition by Clustal Omega. The first and last amino acid of the recombinant pg120 proteins of Subtypes A and B that were used experimentally in this study are shown in bold red and blue letters, respectively.

We used recombinant gp120 proteins purchased from Sino Biological. The gp120 protein of HIV-1 Group M Subtype A (isolate 92RW020, accession # AAT67478.1) consisted of amino acids Glu31-Arg494, and the gp120 protein of Subtype B (isolate BAL, accession # AAA44191.1) consisted of amino acids Glu30-Arg509. Both proteins were expressed in HEK293 cells with a polyhistidine tag at the C-terminus and lyophilized from sterile PBS, pH 7.4 (Table 1). As controls, we used SARS-CoV-2 spike protein S1 subunit (YP_009724390.1; Val16-Arg685) and SARS-CoV-2 spike protein (B.1.1.529/Omicron variant) S1 subunit (YP_009724390.1; Met1-Arg685), polyhistidine-tagged recombinant proteins expressed in HEK293 cells. SARS-CoV-2 spike protein S1 subunit shares low amino acid similarity with the gp120 protein, 7.3% amino acid identity with gp120 of Subtype A and 8.5% with Subtype B, allowing us to test the specificity of the detected responses. Both gp120 of HIV-1 and S1 of SARS-CoV-2 can be cleaved off by furin and may participate in promoting cardiovascular pathologies (Suzuki, 2020; Suresh & Suzuki, 2021).

Human pulmonary artery endothelial cells were treated with recombinant gp120 of Subtypes A and B to determine how these proteins may change the gene expression patterns. We first used R&D Proteome Profiler Array Human XL Oncology Array Kit that allows for simultaneously monitoring the expression of various proteins, many of which are involved in cell signaling regulation.

After treating cells for 20 hours at 1 nM of gp120 proteins as well as SARS-CoV-2 spike proteins as references, cell lysates were prepared and subjected to the protein array. Three separate experiments were performed for each group, the densitometry analysis was performed for each array, and conclusions were made based on a statistical analysis of the results. Each array contained two spots for a protein. We took an average of these two spots to determine the value for the array and performed analyses for three arrays per group.

This approach revealed proteins that were differentially affected by HIV gp120 Subtype A and gp120 Subtype B. One protein whose expression was found to be differentially affected by Subtypes A and B is prostasin/Prss8. Fig. 2A shows the representative images of prostasin protein spots in the protein arrays, demonstrating that cells treated with Subtype B gp120 have higher prostasin expression than those treated with Subtype A gp120. We also show, as a reference, neighboring spots of E-selectin that had similar intensities between Subtypes A and B. The difference between the effects of gp120 of Subtypes A and B on the prostasin expression was found to be statistically significant as indicated in the bar graph (Fig. 2A). On the other hand, the expression levels of E selectin were not significantly different, and the ratio of prostasin to E selectin showed a significant difference between Subtypes A and B.

**Fig. 2:**
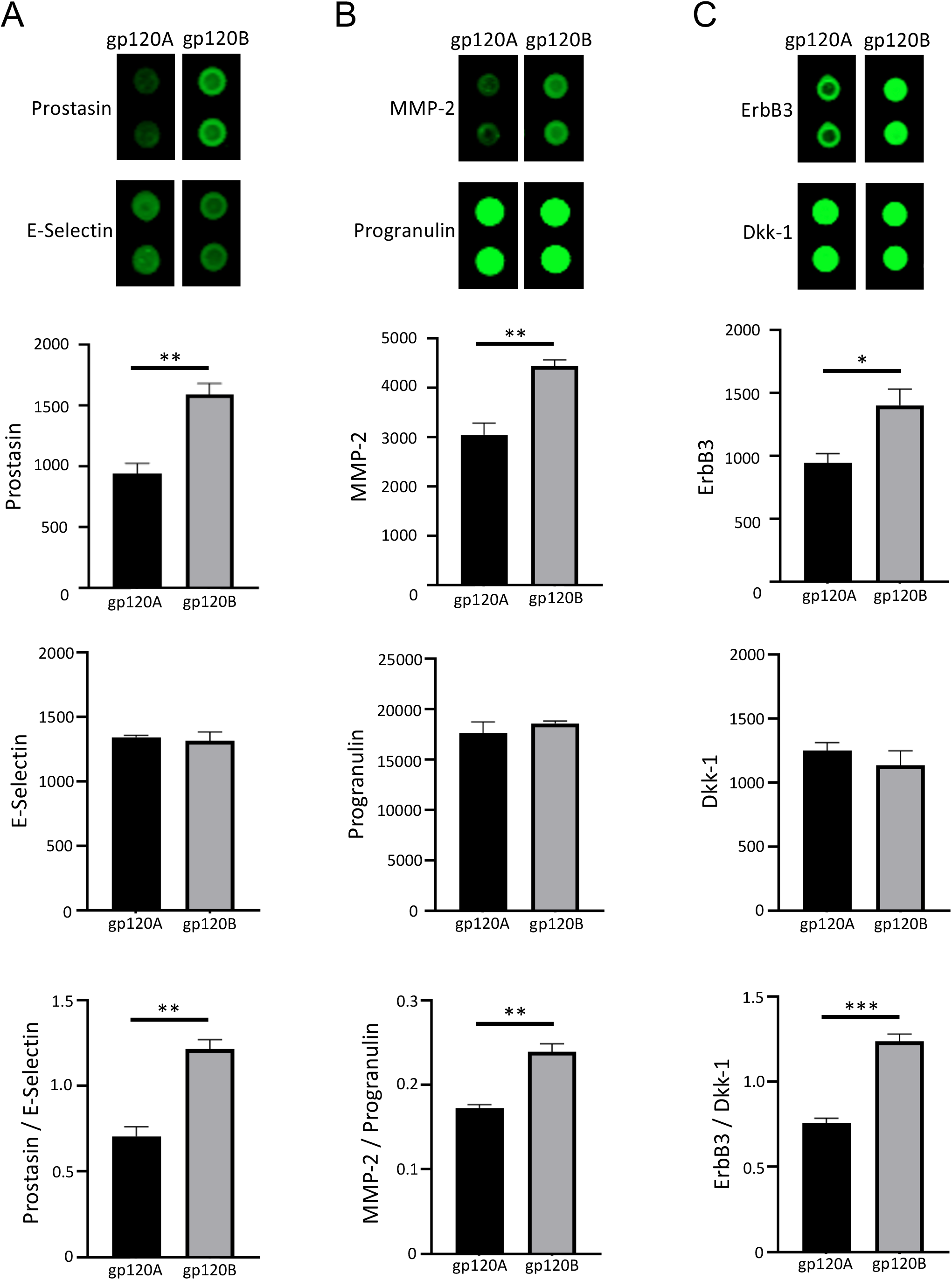
gp120 proteins of HIV-1 Group M Subtype A and Subtype B differentially affect prostasin, MMP-2 and ErbB3. Human pulmonary artery endothelial cells were treated with gp120 of HIV-1 Subtype A (gp120A) or Subtype B (gp120B) at 1 nM for 24 h. Expression patterns of various proteins were monitored using R&D Human XL Oncology Array. (A) Prostasin was found to be higher in gp120B-treated cells, while neighboring E-selectin spots were unchanged. (B) MMP-2 was found to be higher in gp120B-treated cells, while neighboring progranulin spots were unchanged. (C) ErbB3 was found to be higher in gp120B-treated cells, while neighboring Dkk-1 spots were unchanged. Representative images are shown at the top. Densitometry values from two spots from each array were averaged and statistical analysis was performed using results from three separate treatments/arrays. Bar graphs represent means ± SEM (N=3). *P<0.05. **P<0.01. ***P<0.001.

Similarly, matrix metalloproteinase-2 (MMP-2) was found to be differentially expressed after the treatment of cells with either gp120 of Subtype A or Subtype B (Fig. 2B). On the other hand, spots representing progranulin did not exhibit subtype differences, and ratio of these proteins showed significant differences between the effects of Subtypes A and B (Fig. 2B).

The expression of ErbB3 (Her3) was also found to be higher in cells treated with Subtype B gp120 compared to cells treated with Subtype A gp120 (Fig. 2C). The differences in the intensities of these spots were statistically significant. On the hand, the levels of nearby Dkk-1 spots were found to be similar between cells treated with subtypes A and B, and the normalization of ErbB3 values to Dkk-1 values showed a significant difference.

These results revealed, for the first time, that gp120 proteins from different HIV subtypes can exhibit different effects on human host cells.

We further compared the prostasin expression levels in cells treated with gp120 proteins with untreated cells as well as spike protein-treated cells. Fig. 3A shows that gp120 of Subtype A and Subtype B both caused significant downregulation of prostasin compared to untreated cells. This downregulation of prostasin was promoted significantly more potently by Subtype A gp120 compared to Subtype B. Both gp120 subtypes promoted significant downregulation of Eselectin, but to the same extent between the subtypes (Fig. 3B). The analysis of prostasin values normalized with E-selectin demonstrated that only gp120 of Subtype A, but not gp120 of Subtype B, caused a significant downregulation (Fig. 3C). SARS-CoV-2 spike proteins (either originally reported S1 subunit or Omicron S1 subunit) did not have any significant effects on prostasin or on E-selectin expression levels, suggesting that the response for gp120 is specific (Fig. 3).

**Fig. 3:**
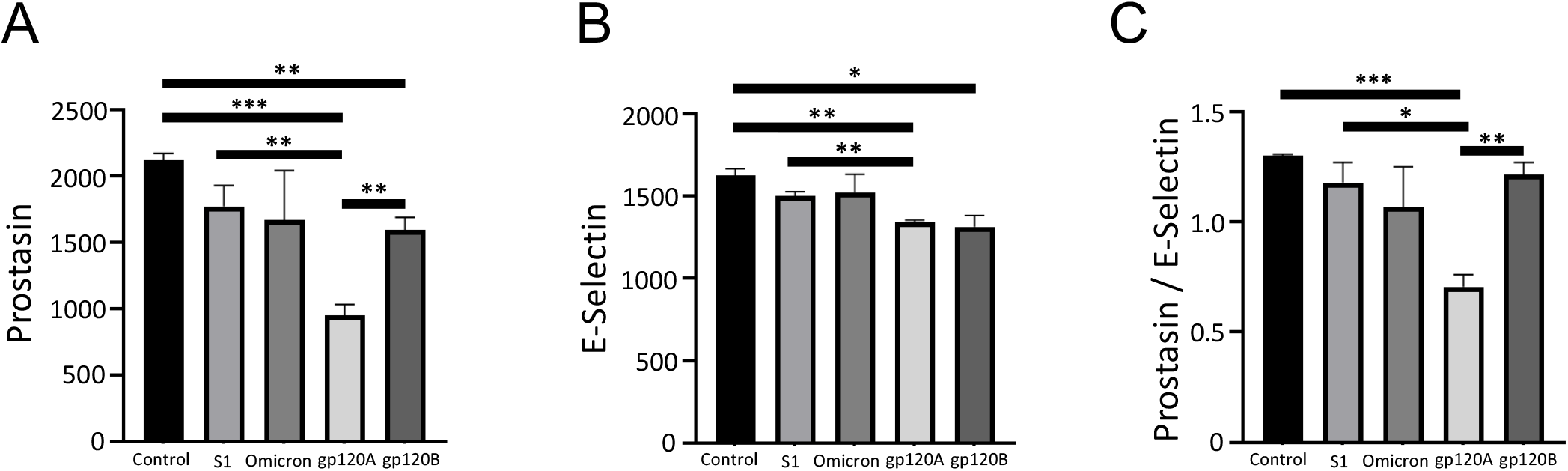
Characteristic of the effects on the prostasin expression. Human pulmonary artery endothelial cells were treated with SARS-CoV-2 spike protein S1, Omicron S1, gp120 of HIV-1 Subtype A (gp120A) or Subtype B (gp120B) at 1 nM for 24 h. Expression patterns of various proteins were monitored using R&D Human XL Oncology Array. Bar graphs represent means ± SEM (N=3) of (A) prostasin expression, (B) E-selectin expression, (C) the ratio of prostasin to E-selectin expression. *P<0.05. **P<0.01. ***P<0.001.

MMP-2 was also found to be significantly downregulated by both Subtypes A and B of gp120 (Fig. 4A). Subtype A was significantly more potent in causing the downregulation of MMP-2. While gp120 proteins of both subtypes also caused significant downregulation of progranulin, no differences were detected in the potencies of the two subtypes (Fig. 4B). The analysis of MMP-2 values normalized with progranulin showed that only Subtype A, but not Subtype B, significantly caused a downregulation (Fig. 4C). We also noted that the spike protein (with the original sequence) caused downregulation of MMP-2 and progranulin (Figs. 4A and 4B). However, the downregulation effects of Subtype A gp120 were found to be more potent than that of the spike protein (Figs. 4A and 4C).

**Fig. 4:**
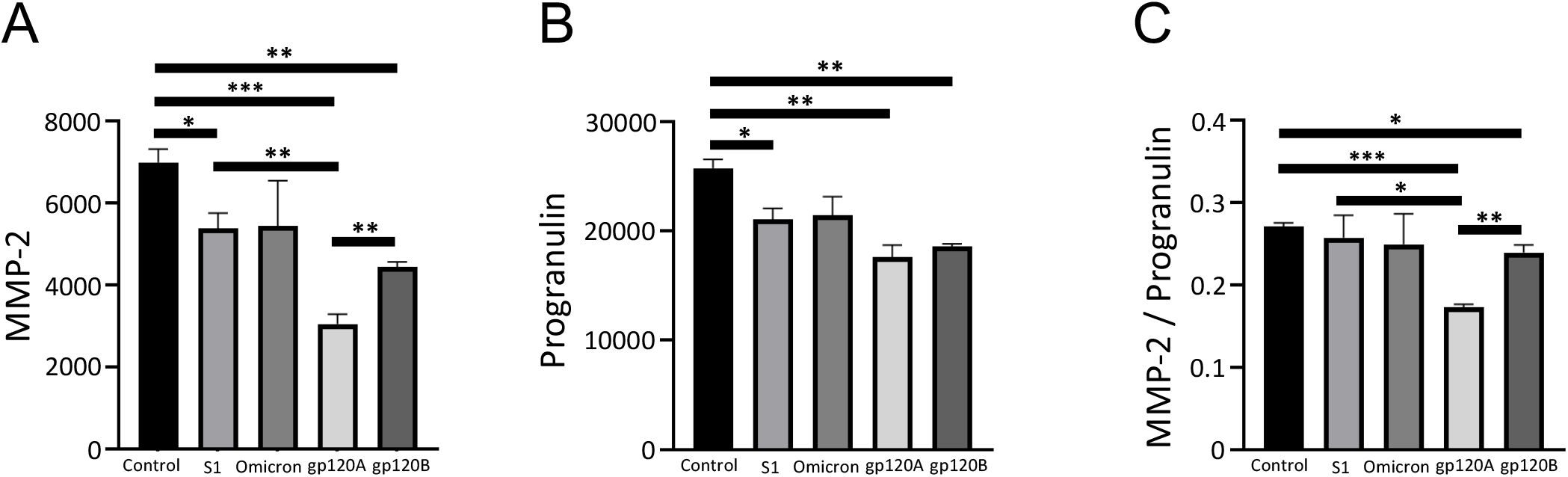
Characteristic of the effects on the MMP-2 expression. Human pulmonary artery endothelial cells were treated with SARS-CoV-2 spike protein S1, Omicron S1, gp120 of HIV-1 Subtype A (gp120A) or Subtype B (gp120B) at 1 nM for 24 h. Expression patterns of various proteins were monitored using R&D Human XL Oncology Array. Bar graphs represent means ± SEM (N=3) of (A) MMP-2 expression, (B) progranulin expression, (C) the ratio of MMP-2 to progranulin expression. *P<0.05. **P<0.01. ***P<0.001.

The analysis of ErbB3 spots revealed that only gp120 of Subtype A, but not of Subtype B, caused significant downregulation of ErbB3 (Fig. 5A). The neighboring protein Dkk-1, that was used as a reference in Fig. 2 as a protein whose expression was not different between cells treated with either Subtype A or B gp120, was found to be significantly downregulated by both subtypes of gp120 (Fig. 5B). Spike proteins did not have any effects of the expression of ErbB3 (Fig. 5A).

**Fig. 5:**
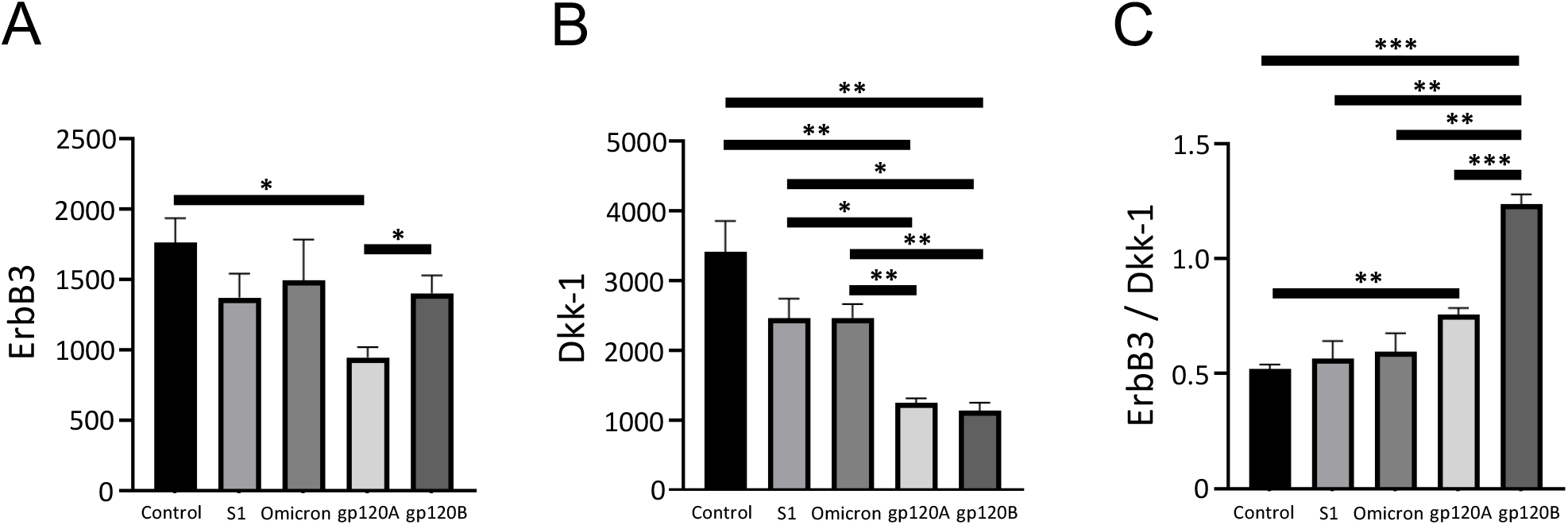
Characteristic of the effects on the ErbB3 expression. Human pulmonary artery endothelial cells were treated with SARS-CoV-2 spike protein S1, Omicron S1, gp120 of HIV-1 Subtype A (gp120A) or Subtype B (gp120B) at 1 nM for 24 h. Expression patterns of various proteins were monitored using R&D Human XL Oncology Array. Bar graphs represent means ± SEM (N=3) of (A) ErbB3 expression, (B) Dkk-1 expression, (C) the ratio of ErbB3 to Dkk-1 expression. *P<0.05. **P<0.01. ***P<0.001.

These results showed that gp120 of HIV Group M Subtype A potently downregulates prostasin, MMP-2, and ErbB/Her3 protein expressions in pulmonary artery endothelial cells, while gp120 of Subtype B has minimal effects.

Conversely, we found that gp120 of Subtype B, but not Subtype A, downregulated monocyte chemotactic protein-2 (MCP-2/CCL8) and MCP-3 (CCL7). Fig. 6 shows that the expression of MCP-2 and MCP-3 are higher in cells treated with gp120 Subtype A compared to cells treated with Subtype B gp120. Quantifications of these experiments determined that these differences were statistically significant (Figs. 6B and 6C). Comparisons with untreated cells revealed that Subtype A gp120 had no effect, while Subtype B significantly downregulated the protein expression of MCP-2. In the case of MCP-3, we found that Subtype A gp120 significantly increases and Subtype B gp120 decreases its expression. Spike proteins did not exert any significant effects.

**Fig. 6:**
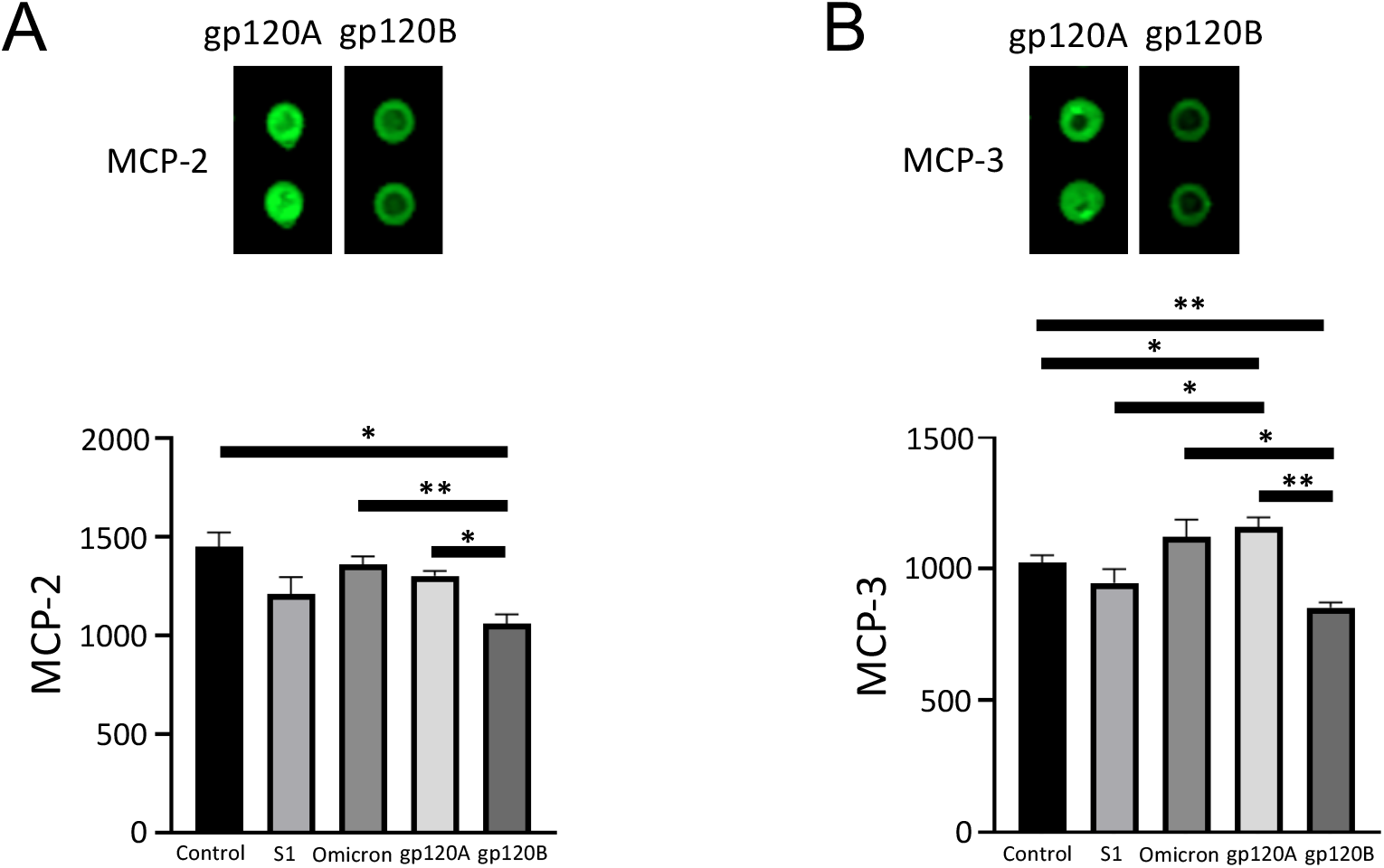
gp120 proteins of HIV-1 Group M Subtype A and Subtype B differentially affect prostasin, MCP-2 and MCP-3. Human pulmonary artery endothelial cells were treated with SARS-CoV-2 spike protein S1, Omicron S1, gp120 of HIV-1 Subtype A (gp120A) or Subtype B (gp120B) at 1 nM for 24 h. Expression patterns of various proteins were monitored using R&D Human XL Oncology Array. (A) MCP-2 was found to be higher in gp120A-treated cells. (B) MMP-3 was found to be higher in gp120A-treated cells. Representative images are shown at the top. Densitometry values from two spots from each array were averaged and statistical analysis was performed using results from three separate treatments/arrays. Bar graphs represent means ± SEM (N=3). *P<0.05. **P<0.01. ***P<0.001.

We further performed the Human XL Cytokine Array analysis and found that thymus- and activation-regulated chemokine (TARC/CCL17) is one protein that is differentially expressed in response to gp120 of Subtypes A and B. As shown in Fig. 7A, the TARC expression is downregulated by gp120 of both subtypes, but Subtype B was significantly more potent than Subtype A. By contrast, in this array, angiopoietin-1 protein expression was found not to change in response to gp120 of either subtype (Fig. 7B). The value of TARC normalized to angiopoietin-1 was significantly lower in cells treated with Subtype B gp120 compared to Subtype A.

**Fig. 7:**
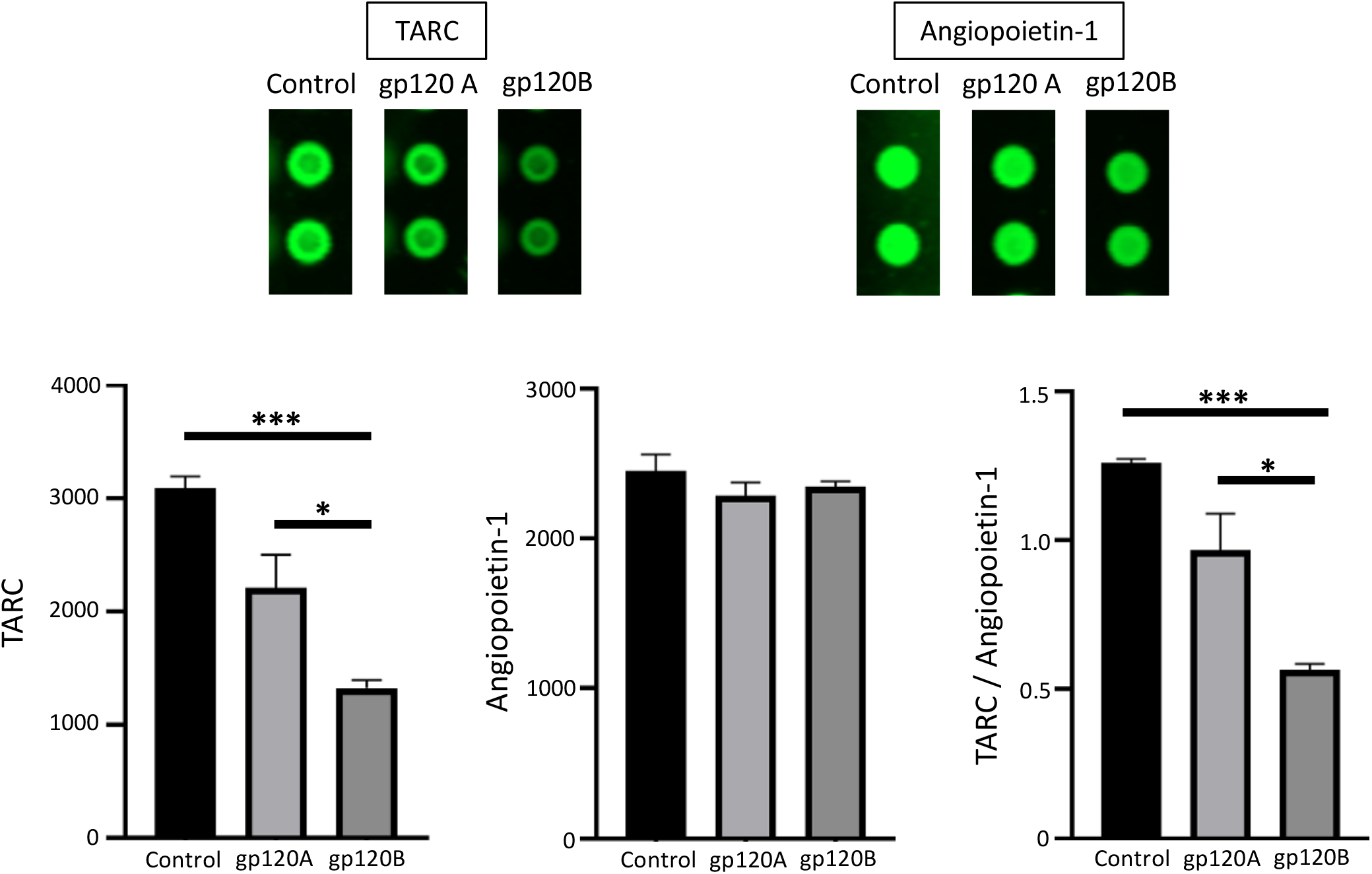
gp120 proteins of HIV-1 Group M Subtype A and Subtype B differentially affect TARC expression. Human pulmonary artery endothelial cells were treated with gp120 of HIV-1 Subtype A (gp120A) or Subtype B (gp120B) at 1 nM for 24 h. Expression patterns of various proteins were monitored using R&D Human XL Cytokine Array. The TARC expression was found to be higher in gp120A-treated cells, while neighboring angiopoietin-1 spots were unchanged. Representative images are shown at the top. Densitometry values from two spots from each array were averaged and statistical analysis was performed using results from three separate treatments/arrays. Bar graphs represent means ± SEM (N=3). *P<0.05. **P<0.01. ***P<0.001.

Taken together, gp120 proteins of HIV-1 Group M Subtypes A and B that share ~70% homology in their amino acid sequences differ in how they affect human host vascular endothelial cells.

## Discussion

The majority of HIV-1 infections in eastern Africa and former Soviet Union countries are caused by Subtype A, while Subtype B is the most prevalent subtype in western countries (Nikolopoulos, 2016; Saad, 2006). As a consequence, much research has been carried out on Subtype B, and it is unknown whether the biological actions of the gp120 of these two subtypes differ.

Advances in ART resulted in the long-term survival of HIV-positive individuals but also increased clinical concerns of complications that affect these individuals such as vascular diseases including PAH. To investigate if there could be differences in vascular complications developed in patients infected with HIV-1 Subtype A and Subtype B, we tested the hypothesis that human host vascular cells may respond differently to gp120 viral fusion protein of HIV-1 Group M Subtype A compared to Subtype B. In HIV, the viral fusion protein, gp120, binds to host cell receptors, particularly the CD4 of T cells, to allow the virus to enter the cell. In other cell types, recombinant HIV gp120 has been shown to activate cell signaling events (Schecter, 2001; Green, 2014; Del Corno, 2014; Yang, 2009; Hioe, 2011).

The amino acid sequence comparison analysis of the gp120 protein of Subtype A and Subtype B revealed these proteins only share ~70% amino acid identity. The application of these proteins to cultured human pulmonary artery endothelial cells in conjunction with the use of Proteome Profiler Arrays revealed that HIV-1 Subtypes A and B elicit different protein expression changes. The major differences between the proteins consist of the absence of the TNGNDTNTTSSSRGMV sequence at 137-152 position in gp120 of Subunit A. Further investigations including the role of this region are needed to determine the molecular mechanism of the differential actions of gp120 of Subtypes A and B.

In summary, these results, for the first time, showed that gp120 of HIV-1 Group M Subtypes A and B exhibit different actions on human host cells, providing the basis for the need for rigorously studying the molecular differences between the actions of the HIV-1 components of different subtypes. Such efforts may offer important insights into developing therapeutic strategies to prevent and/or treat PAH and other vascular complications in HIV-positive individuals in accordance with the HIV subtypes affecting different countries.

## Acknowledgements

This research was funded by the National Institutes of Health (NIH), grant numbers R21AI142649 (to Y.J.S.), R03AG059554 (to Y.J.S.), R03AA026516 (to Y.J.S.), R21AG073919 (to Y.J.S. and S.G.G.), R03AG071596 (to Y.S. and S.G.G.), R01GM124020 (to T.I.B.), and R01CA252969 (to T.I.B.). The content is solely the responsibility of the authors and does not necessarily represent the official views of the NIH.

